# Diverse soil RNA viral communities have the potential to influence grassland ecosystems across multiple trophic levels

**DOI:** 10.1101/2021.06.28.448043

**Authors:** Luke S. Hillary, Evelien M. Adriaenssens, David L. Jones, James E. McDonald

## Abstract

Grassland ecosystems form 30-40%^1^ of total land cover and provide essential ecosystem services, including food production, flood mitigation and carbon storage^2^. Their productivity is closely related to soil microbial communities^3^, yet the role of viruses within these critical ecosystems is currently undercharacterised^4^ and in particular, our knowledge of soil RNA viruses is significantly limited^5^. Here, we applied viromics^6^ to characterise soil RNA viral communities along an altitudinal productivity gradient of peat, managed grassland and coastal soils. We identified 3,462 viral operational taxonomic units (vOTUs) and assessed their spatial distribution, phylogenetic diversity and potential host ranges. Soil types exhibited showed minimal similarity in viral community composition, but with >10-fold more vOTUs shared between managed grassland soils when compared with peat or coastal soils. Phylogenetic analyses of viral sequences predicted broad host ranges including bacteria, plants, fungi, vertebrates and invertebrates, contrasting with soil DNA viromes which are typically dominated by bacteriophages^7^. RNA viral communities therefore likely have the ability to influence soil ecosystems across multiple trophic levels. Our study represents an important step towards the characterisation of terrestrial RNA viral communities and the intricate interactions with their hosts, which will provide a more holistic view of the biology of economically and ecologically important terrestrial ecosystems.

---

Within terrestrial ecosystems, DNA viruses are known to play essential roles in microbial community dynamics and carbon biogeochemical cycling^8–11^, however, the role of RNA viruses is currently understudied. Size-fractionated metagenomes (viromes)^12^ of virus-like-particles (VLP) have been shown to detect 30-fold more sequences of DNA viruses than bulk-soil metagenomes^6^ and viral RNA for use in viromics studies can be readily extracted from water, sewage and sediments^13–15^. Genomes and genome fragments of RNA viruses can be detected in soil metatranscriptomes using the universally conserved RNA-dependent RNA polymerase (RdRP) gene marker^5^ but to date, and to the best of our knowledge, there have been no published studies that apply viromics to the study of soilborne RNA viruses.

In this work, we characterised the soil RNA viromes of five contrasting soil types along a typical temperate oceanic grassland altitudinal productivity gradient^16^ (Figure 1a–d). The top 10 cm of soil was evenly sampled from triplicate 5×5 m grids at each site and VLPs were purified from 100 g of soil from each sample. RNA from purified VLPs was DNase treated and reverse-transcribed to cDNA and used to prepare fifteen libraries plus one RNA virome extraction and one library preparation negative control for high-throughput Illumina sequencing. Filtered sequencing reads were co-assembled and contigs >300 bp were used in further analysis. Genes were predicted by Prodigal^17^ and searched for the RNA viral hallmark gene RNA dependent RNA polymerase (RdRP) using HMMER^18^ and five HMMs built from multiple sequence alignments of the RdRPs from the five major RNA viral phyla^19^.

**Figure 1.**
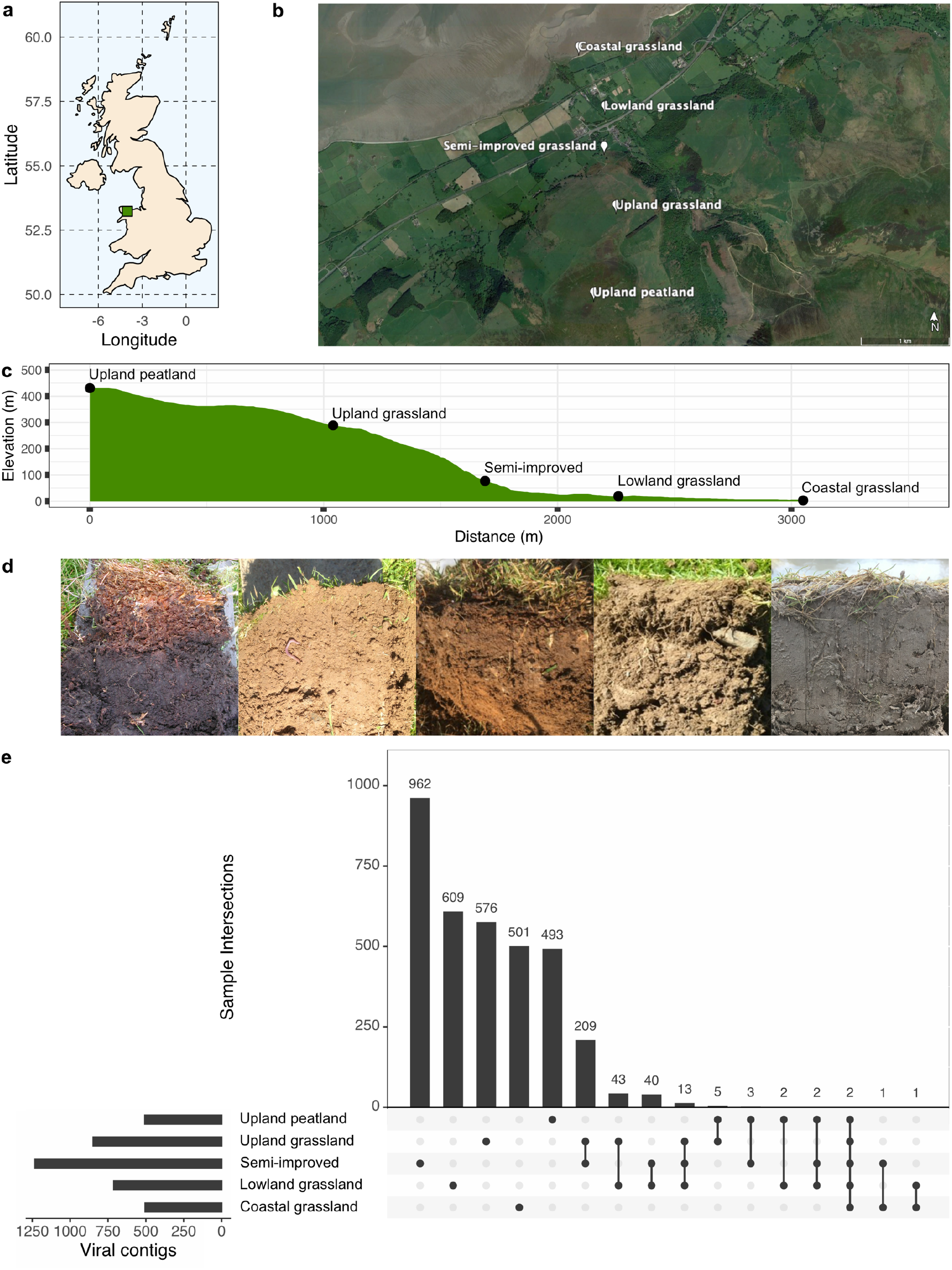
– Soil samples were taken along an altitudinal primary productivity gradient in North Wales, UK (a). Sampling sites included upland peat, three forms of grassland under different management regimes (unimproved upland, semi-improved and improved lowland grassland) and unmanaged coastal grassland (b - made using Google Earth Pro). Elevation varied by 400 m along the transect (c). Site co-ordinates and soil descriptions can be found in supplementary Table 1. Images of soils from each site can be seen in (d) from left to right: upland peatland, upland grassland, semi-imporved grassland, lowland grassland and coastal grassland. An Upset plot of the distribution of identified viral contigs (e) demonstrates how whilst the majority of vOTUs are found solely at each site, the managed grassland sites share more vOTUs in common than with the upland peat or coastal grassland sites, with the coastal grassland site being almost completely unique.

A total of 3,471 contigs that clustered at 95% identity over 85% of the contig length^20^ and contained putative viral RdRP genes were taken forward and considered as viral operational taxonomic units (vOTUs). Read mapping was used to identify vOTUs present within a sample if the horizontal genome coverage was ≥50% (figure 1e). Few vOTUs were shared between sites (0.79-32% per site) with the managed grassland sites showing the most similarity. 97-99% of vOTUs shared by managed grassland sites were shared with at least one other managed grassland site. The coastal grassland site shared the least vOTUs with any other site, whilst the upland peatland site, although markedly different, shared more vOTUs in common with managed grassland sites it was geographically closer to. This could reflect similarities between those habitats, or result from viral particles being transferred between these habitats by ground/ surface water runoff. As the horizontal coverage threshold used to determine vOTU presence can influence the sensitivity and precision of detection^21^, the same analysis was repeated with 25%, 75% and 95% horizontal genome coverage and the same pattern of higher overlap between managed grassland sites than with upland peatland and unmanaged coastal grassland sites was repeated (see supplementary figure 1).

Relative abundance was calculated using TPM (transcripts per kilobase million) values for vOTUs identified as present in each sample and analysed via non-metric multidimensional scaling (NMDS – figure 2). Each site is distinctly separate, with dense collections of vOTUs (grey triangles) located between each sampling replicate, with managed grassland sites positioning closer together than to upland peatland or unmanaged coastal grassland sites. Figures 1e and 2 also show the lack of a clear core soil RNA virome at the vOTU level, and that a combination of soil type, plant coverage and land management may be determining factors of soil RNA viral abundance and diversity.

**Figure 2.**
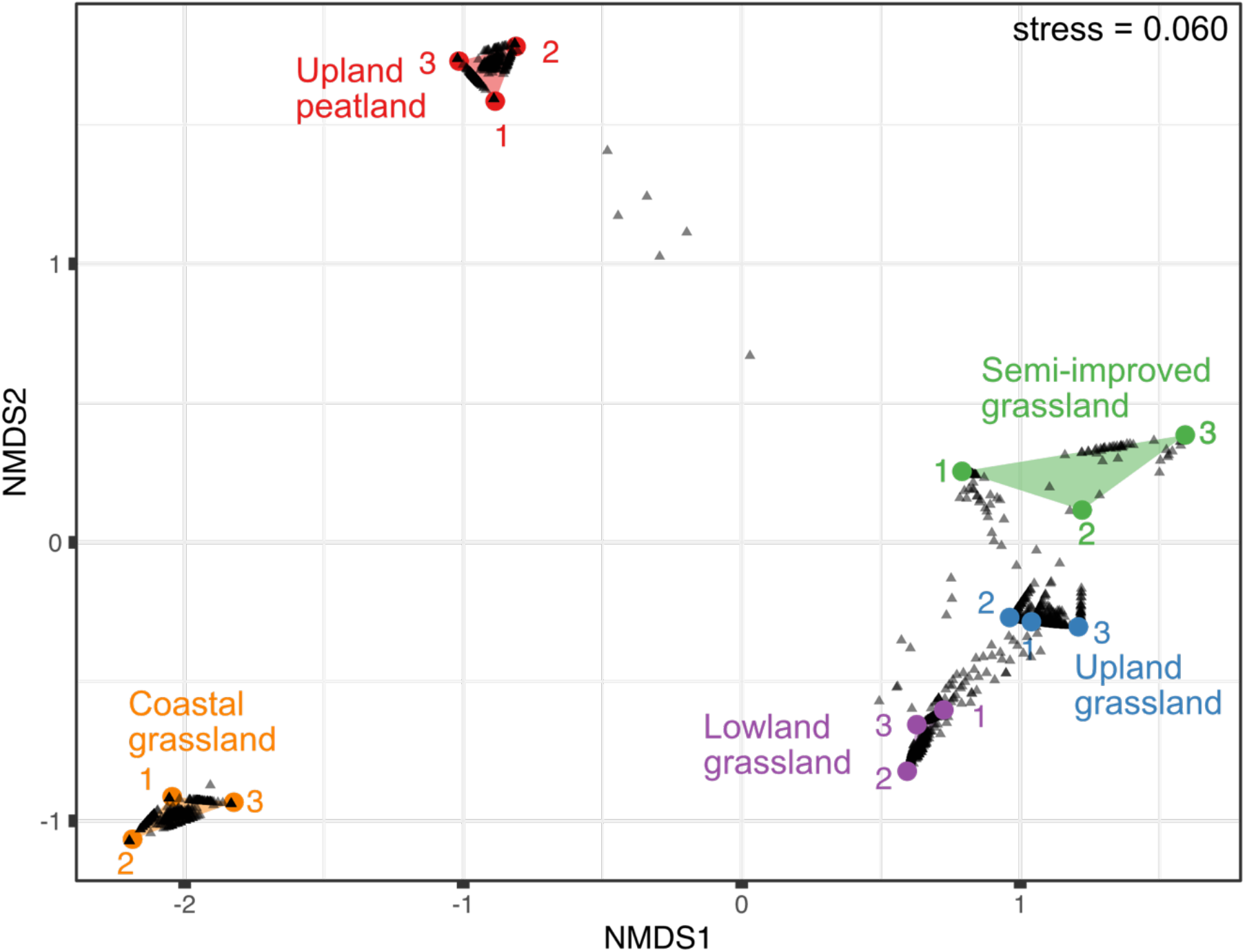
– NMDS of vOTU relative abundance in 5 contrasting soil types along an altitudinal primary productivity gradient. Managed grassland sites cluster closely together whilst peatland and coastal RNA viromes are clearly separated. Whilst a small number of vOTUs (grey triangles) can be seen to be shared between upland-peat and grassland sites, the coastal grassland site is distinctly separate.

To explore broader similarities shared between sites, contigs containing RdRP genes were classified based on the broad phylogenetic scheme constructed by Wolf et al.^19^. This divides the *Riboviria* realm into five branches at the phylum level, based on RdRP amino acid multiple sequence alignments (figure 3a). The coastal grassland site is markedly different from the other sites (figure 3b) and a possible gradient appears to be present: as altitude decreases, the proportion of Branch-1 (*Lenarviricota*) increases, whilst the proportion of Branch-3 (*Kitrinoviricota*) decreases. In contrast, the relative abundance of Branch-2 (*Pisuviricota*) stays broadly similar between each site. The RNA viromes are all heavily dominated by positive-sense single-stranded RNA (+ssRNA) viruses, with the double-stranded RNA (dsRNA) Branch-4 viruses (*Duplornaviricota*) far fewer in relative abundance and mostly observed in the semi-improved and coastal grassland samples. Only four negative-sense (Branch-5 - *Negarnaviricota*) RNA (-ssRNA) vOTUs were identified in the whole study and these were exclusively found in the managed grassland sites. Similar patterns are observable in global RNA viral diversity, where the majority of dsRNA and -ssRNA viruses are nested within the Branch-3 *Kitrinoviricota*^22^.

**Figure 3.**
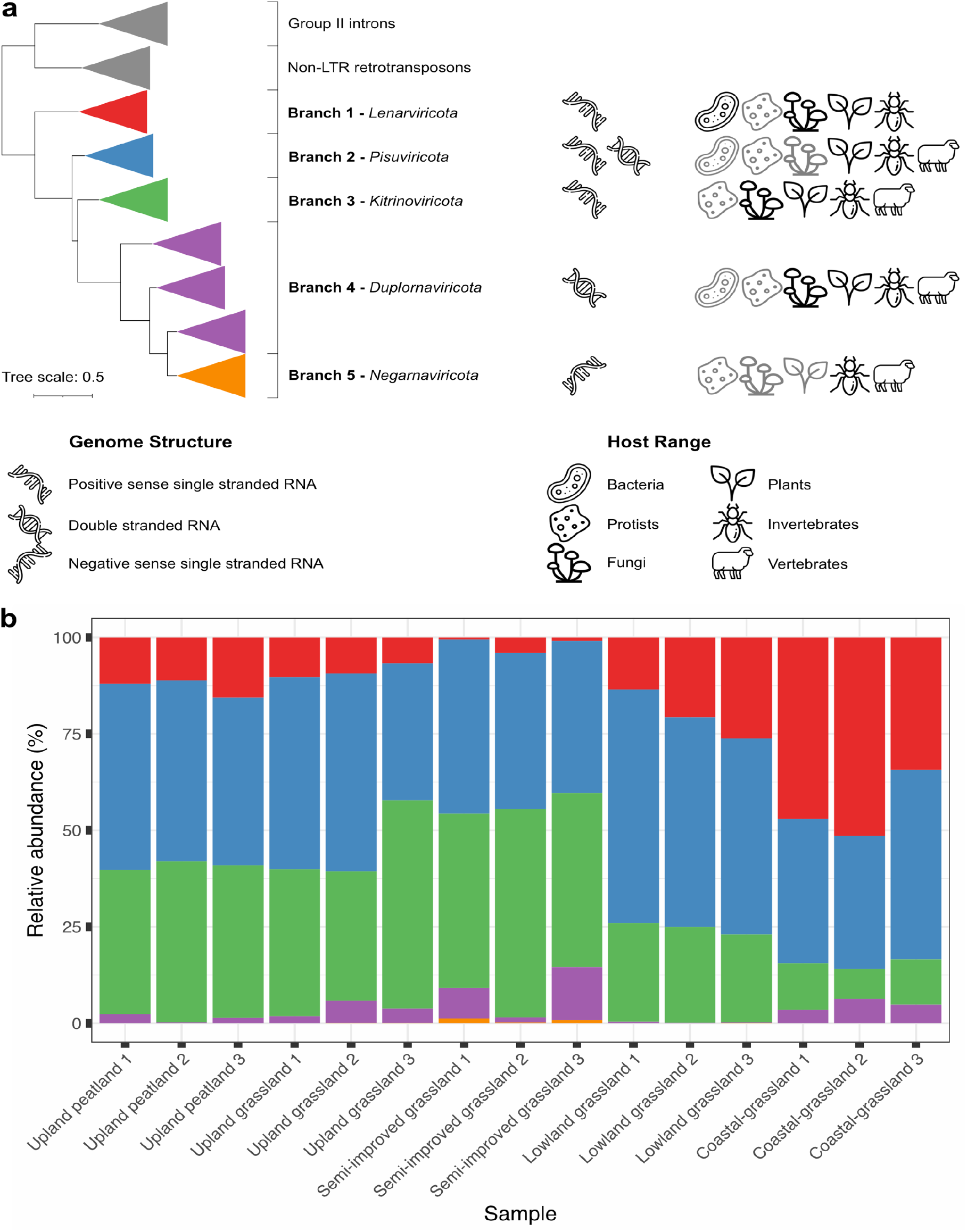
– Phylogenetic organisation, genome structure and host range (a) of the five proposed phyla of RNA viruses, based on RdRP multiple sequence alignments (adapted from Wolf et al.). Grey icons indicate limited known numbers of viruses infecting these hosts. vOTUs from 3 independent samples of 5 contrasting soil types along an altitudinal primary productivity gradient were classified into one of these five branches and relative abundances calculated from TPM values of mapped reads (b). Branch 2 *Pisuviricota* remain broadly similar between samples whilst Branch 1 *Lenarviricota* and Branch 3 *Kitrinoviricota* increase and decrease in proportion respectively moving from upland to coastal sampling sites.

To explore the phylogeny of the viruses discovered in this study further, protein alignments of RdRP genes for viruses in this study, reference viruses from Wolf et al.^23^ and sequences from a recently published bulk soil and leaf litter metatranscriptomics study by Starr et al.^5^ were used to generate phylogenetic trees (figure 4, more detailed trees are found in supplementary figure 2). Many viruses found in this study appear as blocks of closely related viruses containing few reference sequences, similar to the observations of Starr et al.^5^ In other regions of the phylogenetic trees, e.g. *Pisuviricota and Kitrinoviricota* (figure 4b and c), novel viruses are fewer in number and evenly distributed across the known RdRP phylogeny.

**Figure 4.**
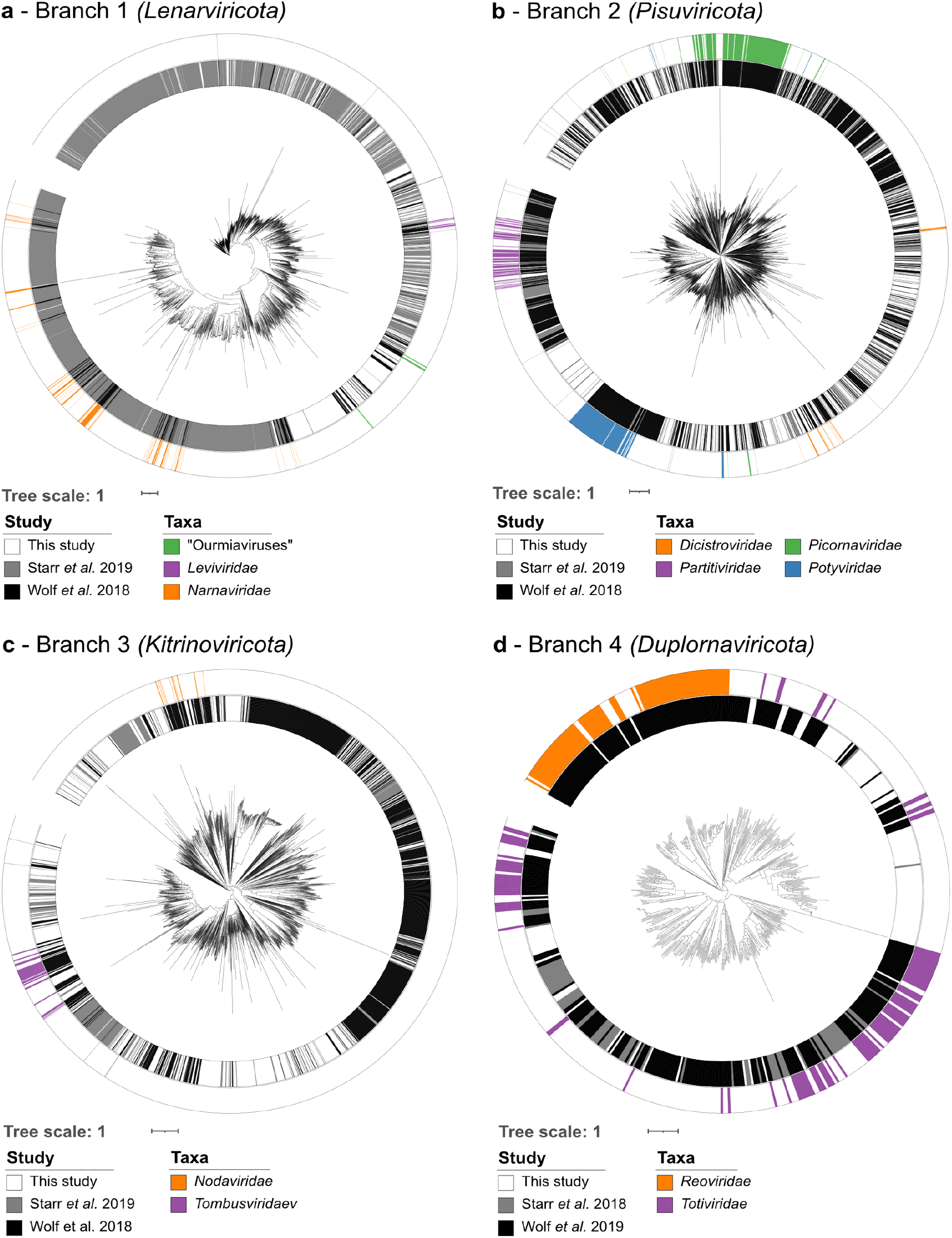
– Phylogenetic trees of RdRP genes based on protein multiple sequence alignments. Sequences found across all soil sites from this study (inner ring, white) were aligned with those used to construct the RNA global taxonomy (Wolf *et al.* inner ring, black) and another soil study (Starr *et al.* inner ring, grey). Global RNA phylogeny is divided into the five proposed RNA viral phyla (a) *Lenarviricota*, (b) *Pisuviricota* (the picornavirus supergroup), (c) *Kitrinoviricota*, (d) *Duplornaviricota*, and *Negarnavirota* (not featured). RdRP genes with established phylogeny of interest are highlighted separately in each panel.

The phylogenetic tree for Branch-1 can be divided into three sections: the class *Leviviricetes*, the family *Narnaviridae* and related mitoviruses, and the newly reclassified family *Botourmiaviridae*^22,24^. Similarly to the work of Starr et al.^5^, this study has detected a large number of potential leviviruses (figure 4a, outer, purple). *Leviviricetes* predominantly infect proteobacteria and their known diversity has recently been significantly expanded^25,26^. We found comparatively few *Narnaviridae* and this may be linked to their structure: *Narnaviridae* members are capsidless +ssRNA viruses that encode no structural proteins and are obligately intracellular^27^. As viromics approaches isolate intact virions, narnaviruses would most likely not be enriched using this technique. In contrast, the encapsidated genus *Ourmiavirs*, within the family *Botourmiaviridae*, comprises plant pathogens with segmented genomes of three ssRNA molecules, each carrying genes for a RdRP, movement protein or capsid protein. 377 ourmia-like vOTUs were identified and are almost exclusively found in managed grassland or upland peat sites (supplementary figure 3), suggesting that this genus of plant viruses may form a larger, more diverse and undercharacterised clade of grassland plant viruses within the *Botourmiaviridae* family. The recovery of complete segmented viral genomes from metagenomics or viromics datasets is particularly challenging due to their segmented nature. Previous studies have used co-occurrence of viral contigs in other publicly available datasets^28^ or sequence homology to known viral species^29^, however the lack of suitable publicly available datasets and extensive horizontal gene transfer can hamper these efforts. *Ourmiaviridae* are known to possess coat proteins that show similarity with highly disparate viruses spanning multiple phyla^19^ and with only three classified species of this genus known, reconstructing their full genomes and classifying novel ourmia-like viruses is particularly challenging. However, their presence here in such high quantities (10% of all detected vOTUs) suggests that they could potentially play an important, but as yet unknown role in grassland ecology. Although viruses are often thought of in terms of pathogenicity, some form persistent mutualistic relationships with their hosts^30^ whilst others have been shown to trigger hypovirulent phenotypes in normally pathogenic plant fungi and are usable as biocontrol agents^31^, creating the prospect that soil viruses may present opportunities for agricultural biotechnology applications.

Branch-2 *Pisuviricota* (Figure 4b), was the most highly represented RNA virus phylum in this study, comprising 40% of identified vOTUs. This is a highly divergent group of viruses with a broad host range and so reliably identifying high taxonomic level hosts for individual vOTUs is challenging. Members of the family *Picornaviridae* infect vertebrates and often cause economically important infections^32^. Relatively few potential *Picornaviridae* were found in this study; however those that were found occupied branches containing various bovine enteric viruses and could be derived from fertiliser manure or sheep dung. Although the separate fields of soil, plant and animal viromics are well established, there are few, if any, studies that consider these separate environments together in order to understand the flow of viruses between them at a community ecology scale.

Of particular interest here are the members of the family *Dicistroviridae* (figure 4b, bottom right, orange). These arthropod-infecting viruses can range from commensal to lethal disease-causing pathogens with significant economic consequences^33^. The dicistro-like viruses found in this study observable in the enlarged tree in Supplementary figure 4a divide into 4 clades: the reference viruses found within the first clade infect crustaceans and the two out of three vOTUs from this study were found in either two or all three coastal samples, the other found in one semi-improved site. The other three clades contain insect-paralysis causing reference-viruses, suggesting that soils may harbour arthropod viruses capable of acutely affecting local mesofauna populations. As soil mesofauna are critical to multiple soil functions^34^, the diversity and ecological roles of arthropod and other invertebrate infecting viruses in soil ecosystems warrants further investigation.

In addition to +ssRNA viruses, viruses belonging to the bisegmented dsRNA *Partitiviridae* family are found within this group, and are capable of infecting plants, fungi and protozoa (see supplementary figure 5). Few vOTUs were found within this group, most likely because, as with *Narnaviridae*, members of the *Partitiviridae* are transmitted exclusively via intracellular mechanisms during spore formation in fungi or ovule/ pollen production in plants^35^. The *Partitiviridae* viruses found in this study may have been released from plant and fungal tissue in the soil during the extraction process or be present as free virions in the extracellular environment. Although this has not been demonstrated empirically here, it is possible to speculate that infection from normally obligate intracellular viruses could occur when mechanical damage occurs to plants and fungi in soils containing infectious and intact virions.

The vOTUs discovered in Branch-3 (*Kitrinoviricota*) are distributed throughout the phylogenetic tree of known RNA viruses and are highly numerate, representing 32.4% of identified vOTUs (figure 4c). Of particular interest here are the three divergent clades in the bottom left quadrant, with the one on the far left containing many known members of the family *Tombusviridae* (blue). These viruses have a wide host range, including plants, protists, invertebrates and vertebrates. The nodaviruses (figure 4c, orange) divide into two categories: alpha-nodaviruses, predominantly isolated from insects but featuring a wide host range under laboratory conditions, and beta-nodaviruses, infecting fish^36^. The noda-like vOTUs identified in this dataset were relatively evenly distributed between the managed grassland and upland peat sites, but none were detected in the coastal grassland (supplemental figure 6).

The phylum *Duplornaviricota* (Branch-4) contain the majority of known dsRNA viruses and comparatively few were detected. *Totiviridae* members (figure 4d - purple) infect fungi, protozoa, vertebrates and invertebrates^19^. vOTUs identified in this study were predominantly found to cluster with isolates that infect animalia (right hand side) but some could be found with the fungi-associated *Totiviridae* (left hand side). Very few vOTUs were found amongst possible *Reoviridae* (figure 4d – orange), with one contig (k127_2512471) showing 97% nucleotide sequence similarity to human rotavirus A (EF554115), found in samples coastal grassland-1 and semi-improved grassland-2.

Only four -ssRNA vOTUs were found in this study and poorly aligned with other known *Negarnaviricota*. -ssRNA virus structure may inhibit detection by viromics: they are almost exclusively lipid-enveloped, occasionally lacking nucleocapsid proteins^37^, and the harsh extraction protocol may lead to virion disruption and loss of viral RNA. This may also be due to the underrepresentation of plant and soil-dwelling arthropod *Negarnaviricota* within nucleotide databases hindering their detection. Our knowledge of this clade is significantly biased towards vertebrate pathogens^38^, however, increased use of high-throughput sequencing has rapidly expanded our breadth of knowledge of *Negarnaviricota* in plants^39^.

Using an altitudinal primary productivity gradient as a source of soils with contrasting ecological properties for RNA virome analysis, this study is the first to apply a direct viromics approach to examine the *in-situ* soil RNA viral community of soil ecosystems. We detected 3,471 vOTUs across five sample sites, and observed site-specific similarities and differences in vOTU relative abundance. The RNA viruses we detected have the potential to infect a range of available hosts, including fungi, bacteria, vertebrates, invertebrates and plants. Therefore, RNA viruses have the potential to influence the grassland soil ecosystem at multiple trophic levels. From a technical standpoint, further development of both wet-lab and bioinformatics techniques are needed to further improve the detection and study of soil RNA viruses. Many RNA viruses have segmented and multipartite genomes, complicating the recovery of full RNA viral genomes from metatranscriptomics and metaviromics datasets. This study found comparatively fewer putative mycoviruses compared to a previous study^5^ examining RdRP containing contigs in soil metatranscriptomics data. This may be due in part to the different structural characteristics and methods of dispersal used by viruses infecting fungi. While it has been shown that viromics outperforms metagenomics in the recovery of DNA viral genomes^6^, the lack of capsid production in key clades of mycoviruses requires consideration when developing future soil RNA viral ecology methodologies.

The impact that soilborne RNA viruses have on their host organisms has only just started to be explored, and future work is needed to establish the many influences they may have on global terrestrial ecosystems. Grassland soil bacterial communities show clear responses to the effects of climate change that are mediated by plant-soil-microbial interactions^40^ and viruses have the potential to influence soil nutrient cycling through host metabolic reprogramming^8^ and their effects on soil microbial community dynamics^5^ similarly to marine viral communities^41^.

Our work demonstrates that RNA viral communities are heavily influenced by location, with upland peatland and unmanaged coastal grassland soils sharing very few vOTUs with managed grassland ecosystems and also showing broad differences at the phylum level. Soilborne RNA viruses identified in this study potentially infect hosts across a wide range of trophic levels and can therefore influence soil ecosystems at a variety of scales. Linking these effects of soilborne RNA virus-host interactions with naturally occurring and anthropogenic environmental processes, will be critical in developing a complete picture of how soil ecosystems respond to environmental change.

## Methods

### Field site description, soil sampling and processing

Five sites along an altitudinal gradient at Henfaes Research Centre, Abergyngregyn, Wales were sampled on 31^st^ October 2018. Three adjacent 5×5 m plots were marked out at each site and approximately 2 kg of soil was extracted from each site between 0-10 cm depth using a 3 cm diameter screw auger with evenly spaced sampling within each grid. The augur was cleaned with disinfectant and a dummy soil core taken and discarded outside of the sampling area prior to sampling each plot. Soil from each plot was sieved separately to 2 mm and stored in 100 g aliquots at −80 °C prior to RNA extraction.

### Viral RNA enrichment and extraction

Virus-like particle extraction was based on protocols developed by Trubl *et al.*^42^ and Adriaenssens *et al.*^13^. A total of 16 samples, three per site and one 100 mL PCR-grade water negative control extraction were processed separately. 100 g of soil per sample was thawed and evenly divided into eight 50 mL centrifuge tubes and suspended in 37.5 mL of amended potassium citrate buffer (1% potassium citrate, 10% phosphate buffered saline (PBS), 5 mM ethylenediaminetetraacetic acid (EDTA), and 150 mM magnesium sulphate (MgSO_4_)). After brief manual shaking, each tube was subjected to 30 seconds shaking followed by 60 seconds vortexing at maximum speed. After physical disruption, tubes were placed on ice on an orbital shaker and shaken at 300 rpm for 30 minutes. Samples were centrifuged for 30 minutes at 3,000 × g, 4 °C. Supernatants were removed to new centrifuge tubes and polyethylene glycol, (PEG - 6,000 MW) and sodium chloride (NaCl) were added to 15% (w/v) and 2% (w/v) respectively to precipitate virus-like-particles overnight at 4 °C. Samples were then centrifuged for 80 minutes at 2,500 × g, 4 °C and pellets resuspended in 10 mL Tris buffer (10 mM Tris-HCl, 10 mM MgSO_4_, 150 mM NaCl, pH 7.5). Samples were filtered through 0.22 μm filters and concentrated to <600 μL using Amicon Ultra-15 centrifugal filter units (50 KDa MWCO, Merck) prior to RNA extraction.

All RNA extraction protocols were used according to the manufacturer’s instructions except where specified. Nucleic acids were extracted using the AllPrep PowerViral DNA/RNA extraction kit (Qiagen) with the addition of 10 μL/ mL 2-β-mercaptoethanol. Co-purified DNA was DNase digested using the Turbo DNA Free kit (Thermo Fisher) using two sequential 30-minute incubations at 37 °C, each using 1 U of Turbo DNase. DNase was inactivated and removed using the supplied DNase inactivation resin and RNA was further purified using the RNA min-elute kit (Qiagen).

### Library preparation, sequencing and initial short read QC

RNA-seq libraries, including an additional library-preparation negative control using an input of PCR grade water, were prepared at the Centre for Genomics Research (CGR), University of Liverpool, using the NEB-Next Unidirectional RNASeq library kit (New England Biolabs). Libraries were pooled and sequenced (150 bp paired end) on one lane of a HiSeq 4000. Initial demultiplexing and quality control performed by CGR removed Illumina adapters using Cutadapt version 1.2.1^43^ with option −O 3 and Sickle version 1.200^44^ with a minimum quality score of 20.

### RNA Virome Data Analysis

Short reads were filtered for read length (<35), GC percentage (<5% and >95%) and mean quality score (<25) using Prinseq-lite v0.20.4^45^. Ribosomal reads were removed using SortMeRNA v3.0.3^46^ using default parameters. Reads from each library were pooled, error corrected using tadpole.sh (mode=correct ecc=t prefilter=2) and deduplicated with clumpify.sh (dedupe subs=0 passes=2) from the BBTools package (v37.76: sourceforge.net/projects/bbmap/). Reads were then co-assembled using MEGAHIT 1.1.3^47^ (-- k-min 27, --k-max 127, --k-step 10, --min-count 1).

### Identification and abundance of viral sequences

Assembled contigs were compared to the NCBI nr complete database (downloaded on 27^th^ November 2019) using Diamond BLASTx^48^ (--sensitive, --max-target-seqs 15, --evalue 0.00001) and taxonomic assignments made using MEGAN v6^49^. All contigs with hits matching cellular organisms or dsDNA/ ssDNA viruses were excluded from subsequent analysis.

HMMs used in RdRP detection were generated from alignments previously published by Wolf *et al*.^19^. Protein coding genes in contigs >300 bp in length were predicted using Prodigal v2.6.3 (-p meta)^17^ and searched for RdRP genes using HMMSearch^18^. Hits with E-values <0.001 and scores >50 were clustered with CD-Hit^50^ to 95% average nucleotide identity across 85% alignment fraction^20^. Each contig was assigned a broad taxonomic classification based on HMMsearch results. Contigs with hits from more than one RdRP branch were assigned to the classification with the lowest E-value.

Reads were mapped to contigs using BBwrap (https://sourceforge.net/projects/bbmap/) and contigs with any mapped reads from either of the two negative control libraries were excluded from further analysis. vOTUs with a horizontal genome coverage of >50% were determined as present for each sample. Any vOTU with coverage of <50% had its abundance reset to zero. Fragments per kilobase million (FPKM) values calculated by BBwrap were converted to TPM using the fpkm2tpm function from the R package RNAontheBENCH^51^. Community analysis was performed using R and the Vegan package^52^. Additional visualisation used the packages ggplot2 and upsetR^53,54^

### Phylogenetic analysis

RdRP sequences from contigs produced by Starr *et al.*^5^ were identified and processed by the same methods described above. These were pooled with those identified by this study and those identified by Wolf *et al*.^19^. Sequences were then aligned using MAFFT^55^ and trees generated with FastTree 2^56^. Trees were visualised with iToL^57^ and annotated with the aid of table2itol (https://github.com/mgoeker/table2itol).

## Author Contributions

LSH, EMA, DLJ and JMD conceived the study and acquired funding. LSH carried out the sample collection and viral RNA extraction and led the data analysis and preparation of the initial draft of the manuscript. All authors contributed to the final version of the article.

## Acknowledgements

This work was supported by funding from the NERC Biomolecular Analysis Facility pilot project competition (project NBAF1158). LSH was supported by a Soils Training and Research Studentship (STARS) grant from the Biotechnology and Biological Sciences Research Council (BBSRC) and Natural Environment Research Council (NE/M009106/1). RNA sequencing library preparation and data acquisition was carried out by the Centre for Genomics Research at the University of Liverpool. Data analysis utilised high performance computing resources from Supercomputing Wales. The authors would like to thank David Fidler for assistance during sample collection, Mike Grimwade-Mann, Sam Morley and Tom Regan for assistance during sample processing, and Dave Chadwick and Robert Griffiths for comments/ advice on data analysis and visualisation.

