## Supplementary Information for "Diverse soil RNA viral communities have the potential to influence grassland ecosystems across multiple trophic levels"

**Supplementary Table 1** – Sampling site descriptions.

| | Site | Location | Soil classification | Soil description | Altitude (m asl) | pH | Electrical conductivity (μs cm$^{-1}$) | Total carbon (%) |
|---|---|---|---|---|---|---|---|---|
| A | Upland peatland | 53° 13' 1.22" N 4° 1' 8.78" W | Non-calcaric lithosol | Very acid upland soil with a wet highly organic topsoil | 431 | 4.27 | 39 | 29.1 |
| B | Upland grassland | 53°13'33.00" N 4° 0' 54.86" W | Typical podzolic brown soil | Freely draining acid loamy soil over rock | 289 | 5.89 | 30 | 11.3 |
| C | Semi-improved grassland | 53° 13' 55.24" N 4° 1’ 2.22” W | Typical podzolic brown soil | Freely draining slightly acid loamy soil | 77 | 4.61 | 27 | 11.2 |
| D | Lowland grassland | 53° 14' 10.98" N 4° 1' 1.74" W | Typical orthic brown soil | Sandy clay loam, freely draining sheep-grazed soil | 19 | 5.78 | 42 | 3.62 |
| E | Coastal grassland | 53° 14' 34.17" N 4° 1' 18.78" W | Saline alluvial gley soil | Silt-textured, poorly draining soil with periodic tidal inundation | 3 | 8.03 | 1810 | 2.89 |

Soils were classified according to Avery (1990). The major properties of the sites and soils are shown in Table S1 above, while a general description of the catena sequence is provided in Farrell et al. (2014) and Withers et al. (2020). The altitudinal gradient represents a primary productivity gradient with more intensive agricultural production at Site D which receives regular fertiliser and lime applications. The mean annual temperature at the bottom and top sites was 10.2 and 7.3 °C respectively, while the gradient in annual rainfall was 1065 to 1690 mm, respectively. All sites had a different vegetation cover (all dominated by grasses) and were grazed by Welsh mountain sheep (*Ovis aries* L.). Soil pH and electrical conductivity were measured in1:2.5 (w/v) soil-to-distilled water extracts using standard electrodes. Total C and N were determined on a TruSpec CN analyser (Leco Corp., St Joseph, MI). Site E is a soil developed on recent marine deposits and contains $CaCO_3$ from shell deposits. Its pedogenic age is ca. 500 years. All other sites have a pedogenic age of ca. 10,000 years. Site A is developed on rhyolite, Sites B and C on

Ordovician schist and shale, and Site D on mixed glacial till. Sites D and E rarely undergo freezing, while Sites A-C experience periodic freezing in winter with winter snow cover often present at Site A. The vegetation at Site A comprises *Festuca ovina* L., *Juncus effusus* L. and *Trichophorum cespitosum* (L.) Hartman. The vegetation at Site B is dominated by *Agrostis canina* L., *Agrostis capillaris* L., *Anthoxanthum odoratum* L. and *Potentilla erecta* (L.) Rauschel. The vegetation at Site C is dominated by *Festuca ovina* L. and *Pteridium aquilinum* L.. Site D is dominated by *Lolium perenne* L. and *Trifolium repens* L. while the vegetation at site E is dominated by *Plantago maritima*, *Festuca* sp. and *Salicornia europaea.*

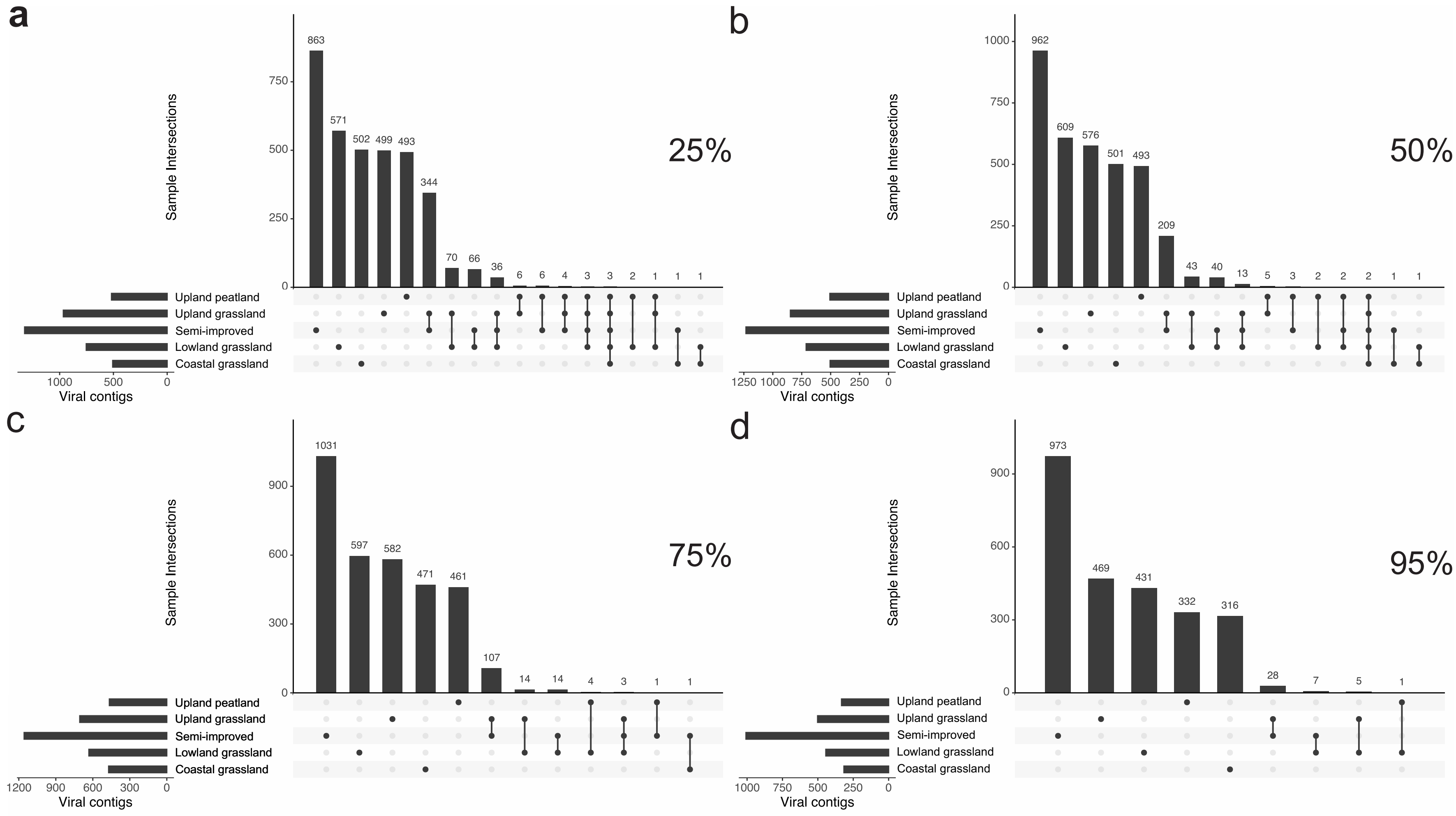


**Supplementary Figure 1** – Comparison of co-occurring vOTUs at (a) 25%, (b) 50%, (c) 75% and (d) 95% horizontal genome coverage thresholds. Similar conclusions on vOTU sharing between sampling sites when varying the alignment fraction required to judge a vOTU present within a sample. vOTUs are most commonly found within one site with the grassland sites sharing more common vOTUs than other habitats. The costal grassland site consistently shared the least number of vOTUs with the other habitats.

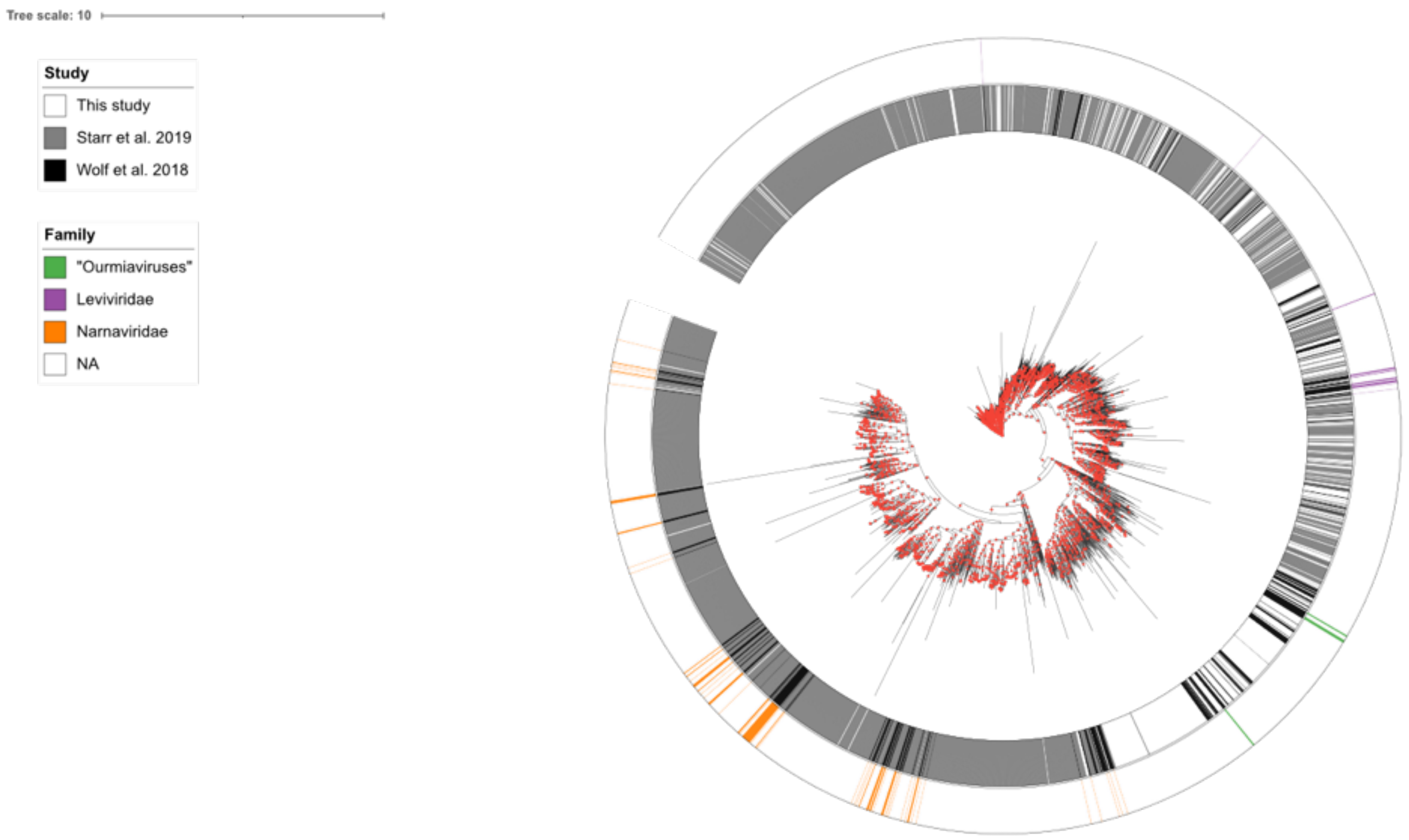


(a) Lenarviricota

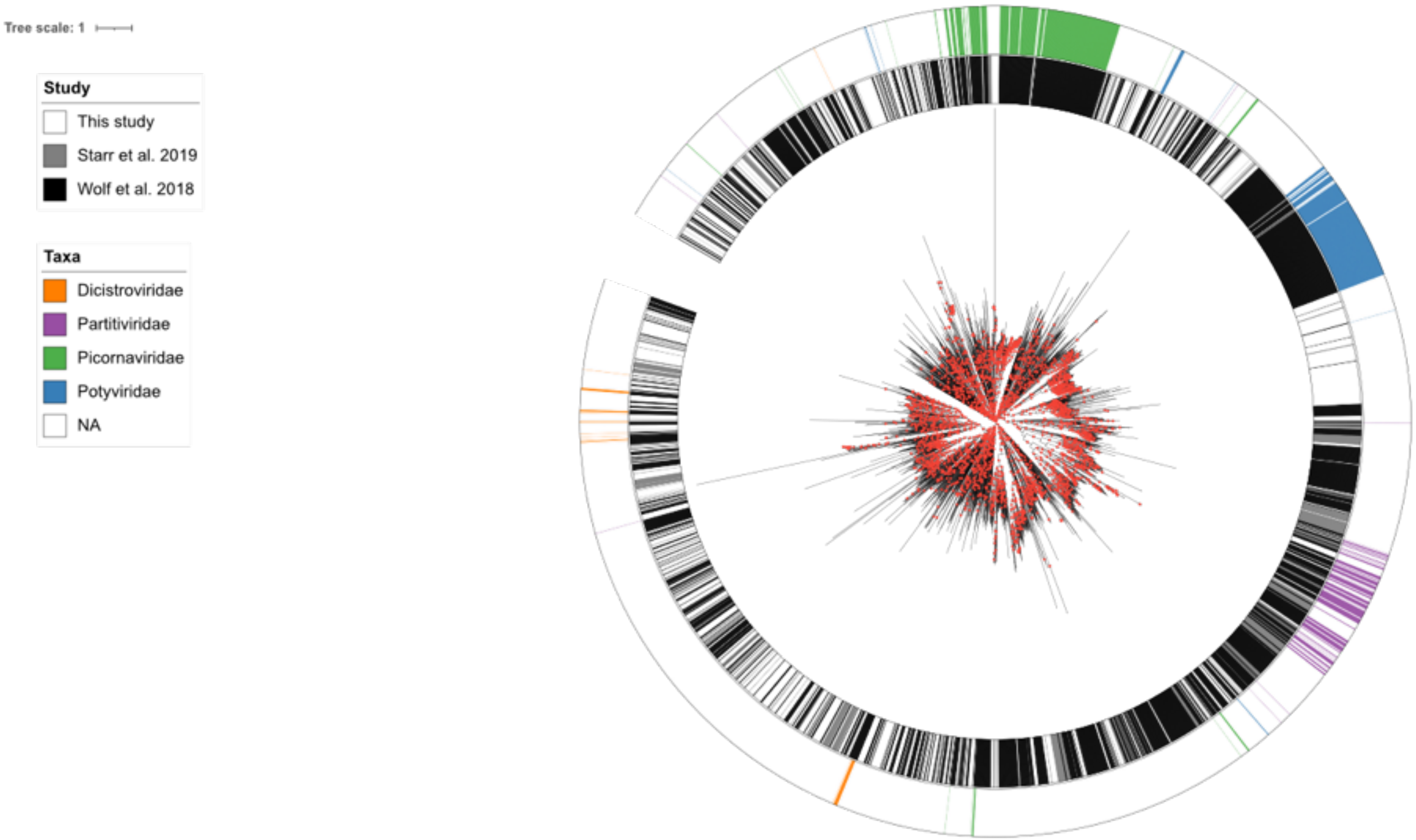


(b) Pisuviricota

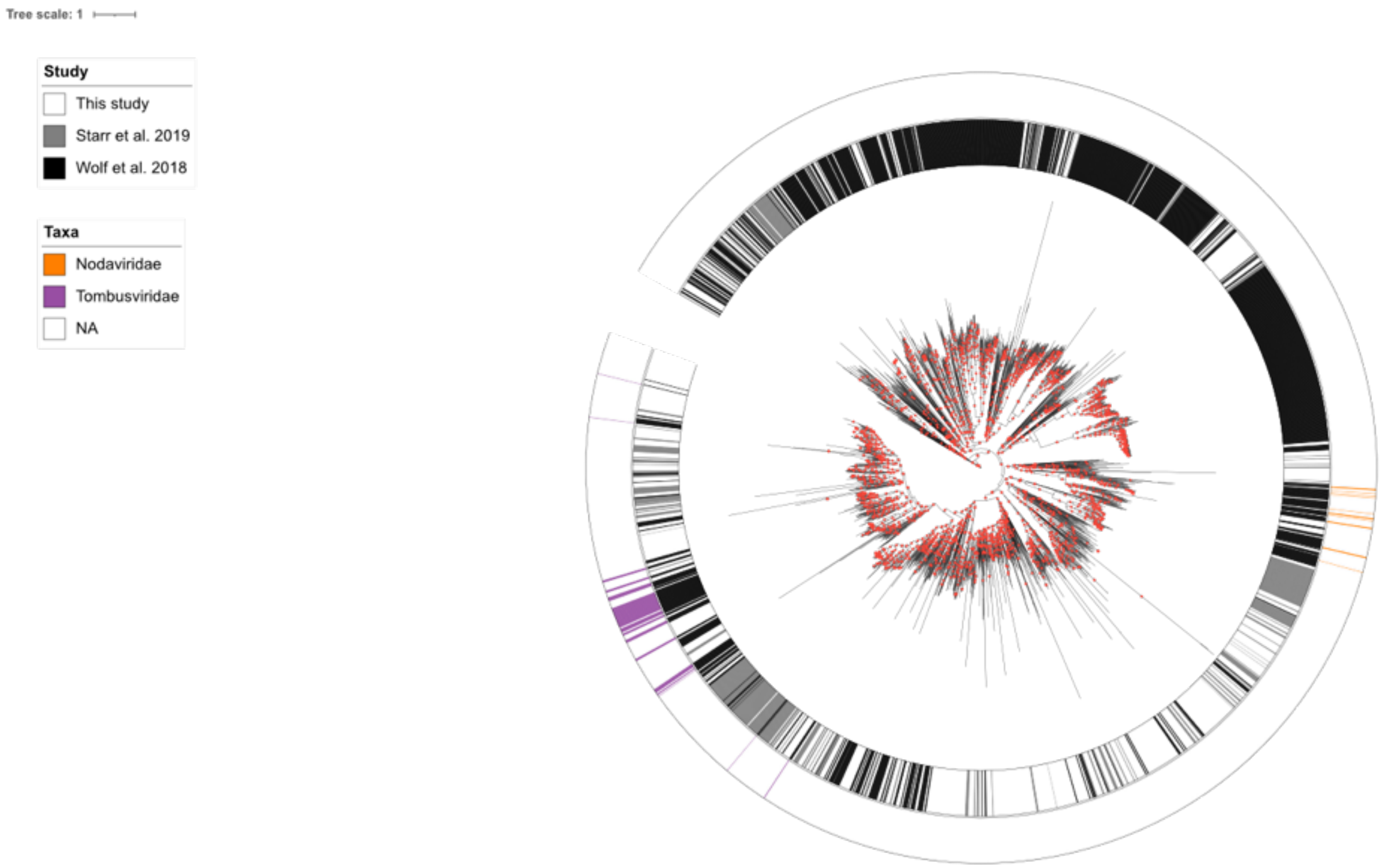


(c) Kitrinoviricota

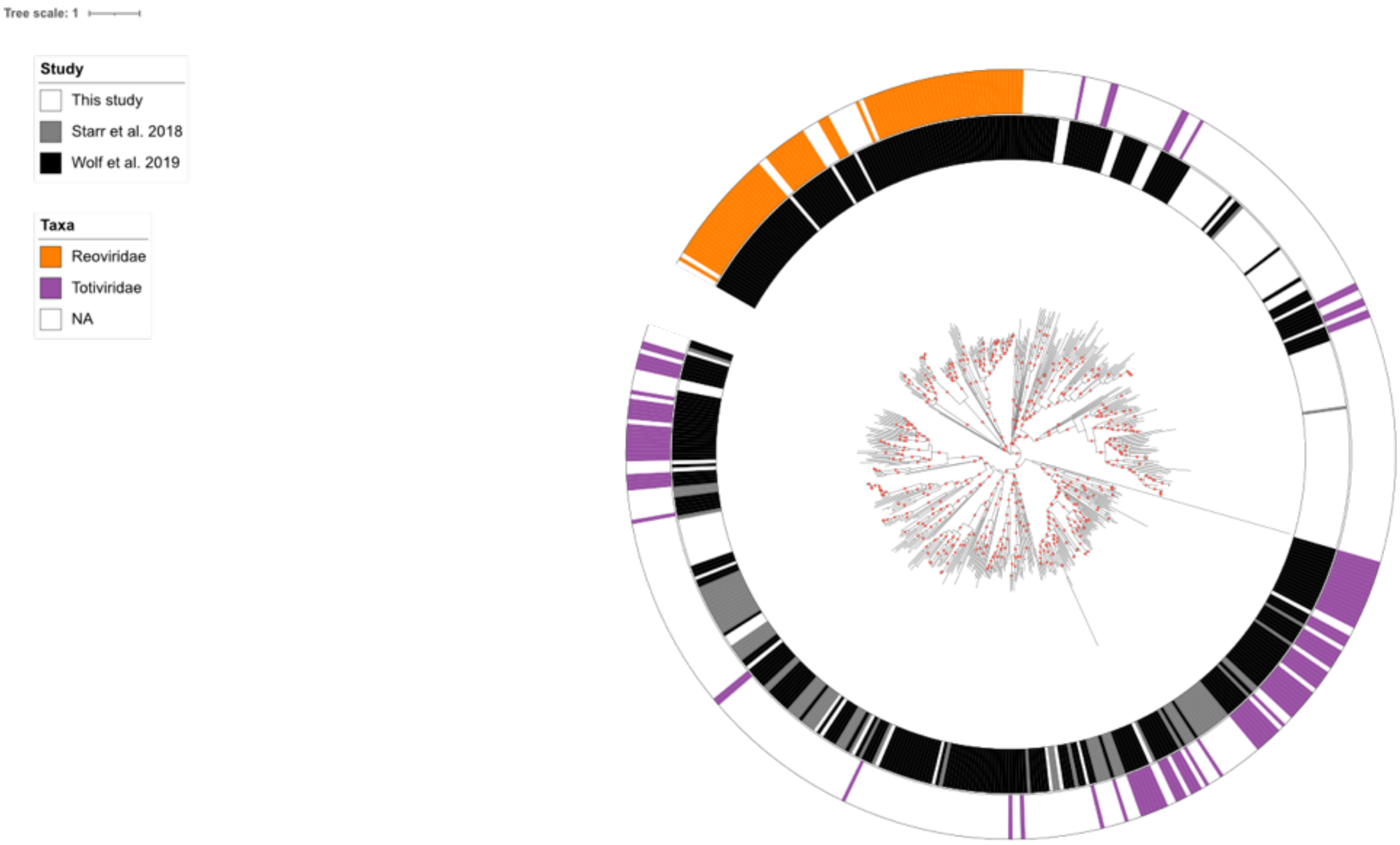


(d) Duplornaviricota

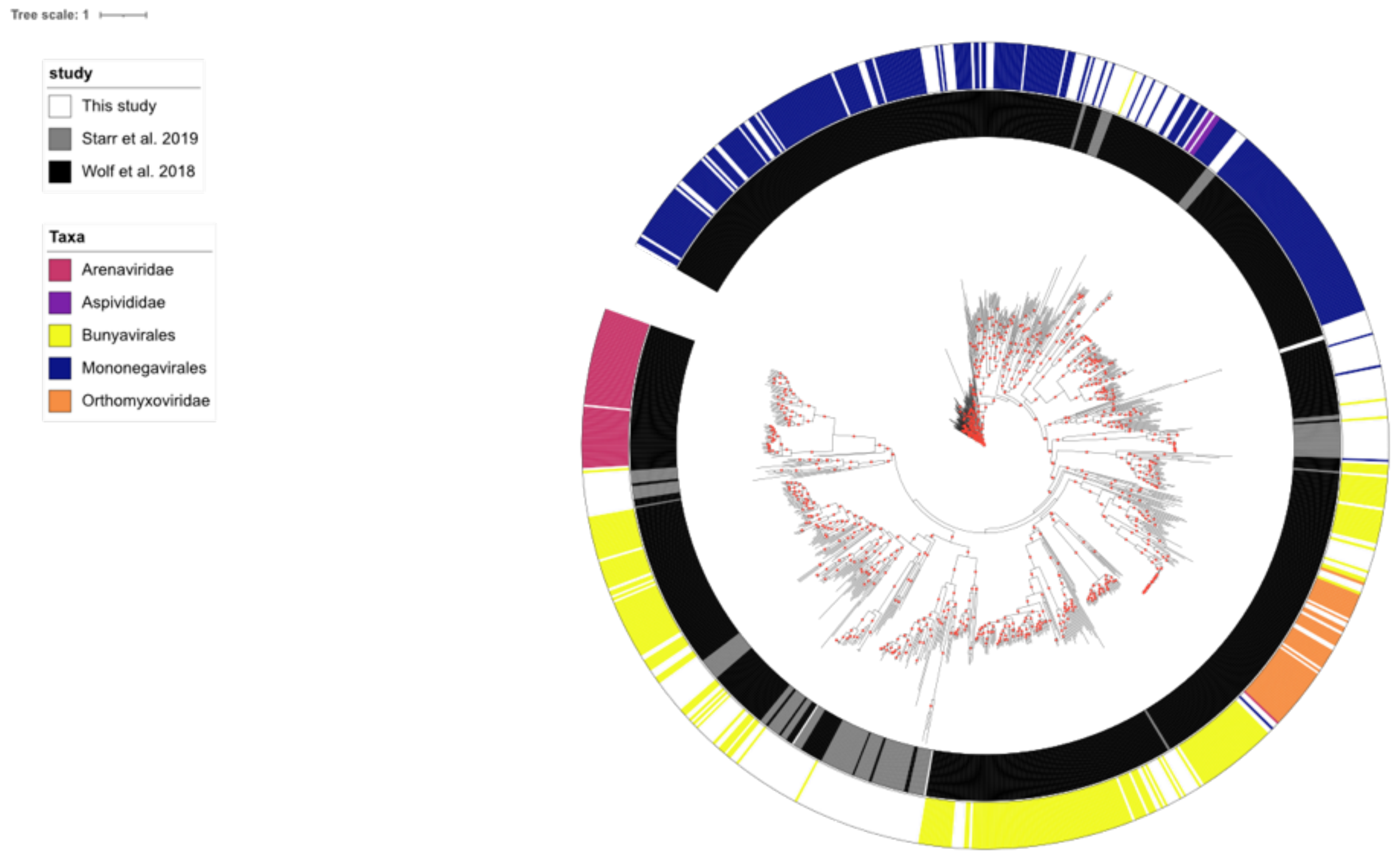


(e) Negarnaviricota

**Supplementary Figure 2** – enlarged phylogenetic trees corresponding to those shown in figure 4 in the main mansucript. Branches with branch support ≥0.6 are indicated by red circles.

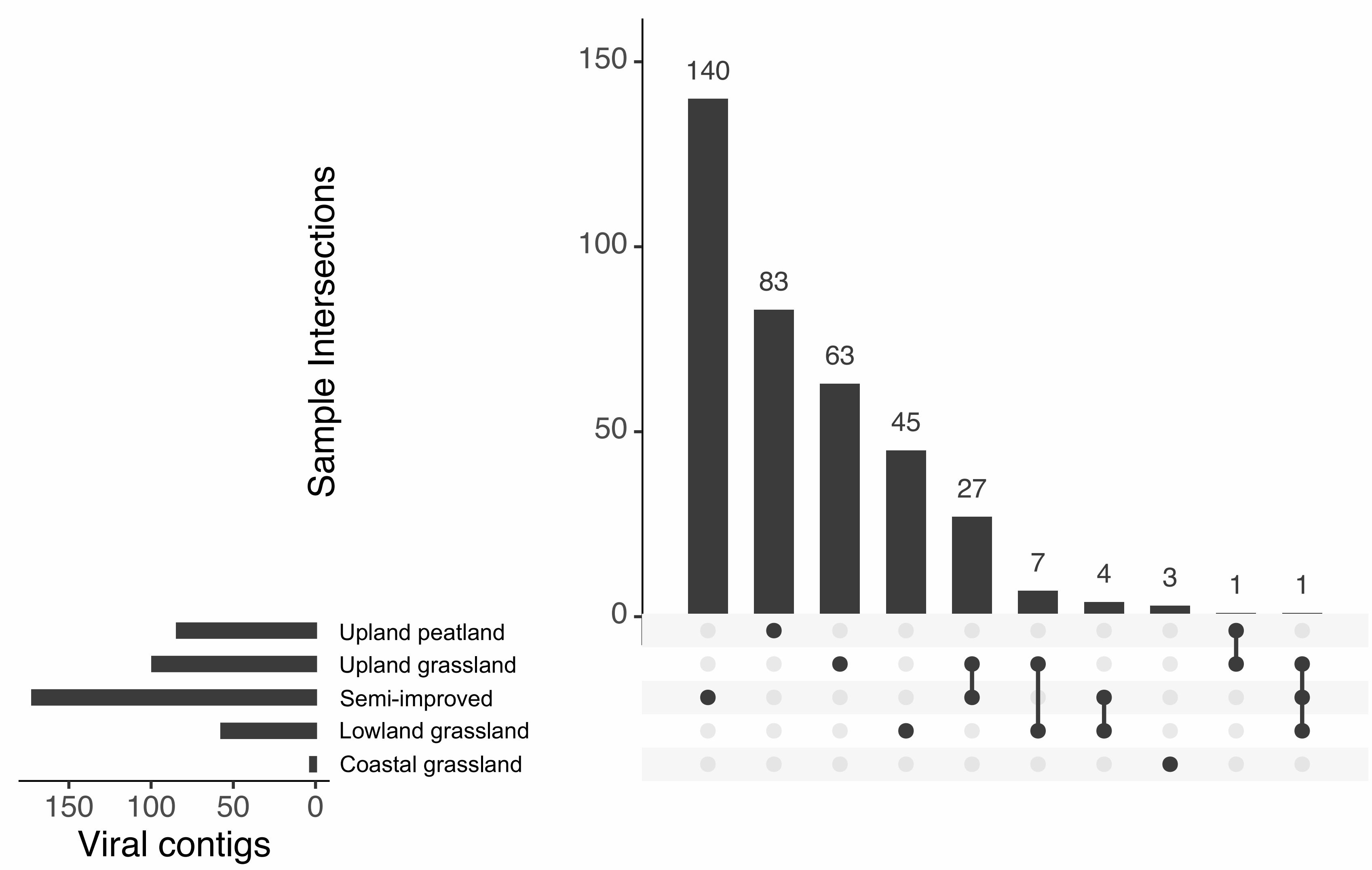


**Supplementary Figure 3** – distribution of putative ourmia-like viruses between sites. Within this study, these viruses are predominantly found in upland grassland/ peatland soils.

(a)

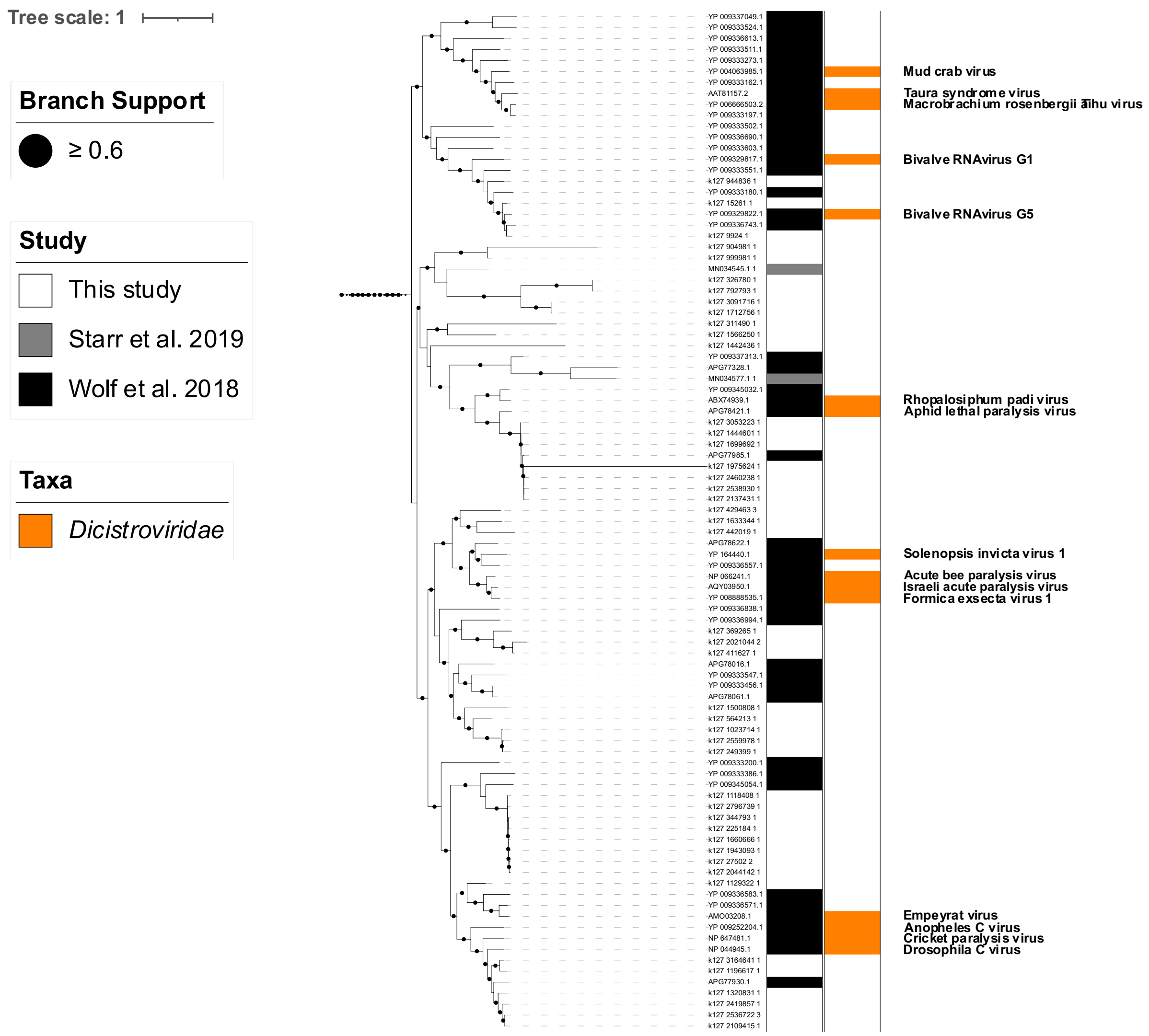
Tree scale: 1
Branch Support
≥ 0.6
Study
This study
Starr et al. 2019
Wolf et al. 2018
Taxa
Dicistroviridae
Mud crab virus
Taura syndrome virus
Macrobrachium rosenbergii Taihu virus
Bivalve RNAvirus G1
Bivalve RNAvirus G5
Rhopalosiphum padi virus
Aphid lethal paralysis virus
Solenopsis invicta virus 1
Acute bee paralysis virus
Israeli acute paralysis virus
Formica exsecta virus 1
Empeyrat virus
Anopheles C virus
Cricket paralysis virus
Drosophila C virus

(b)

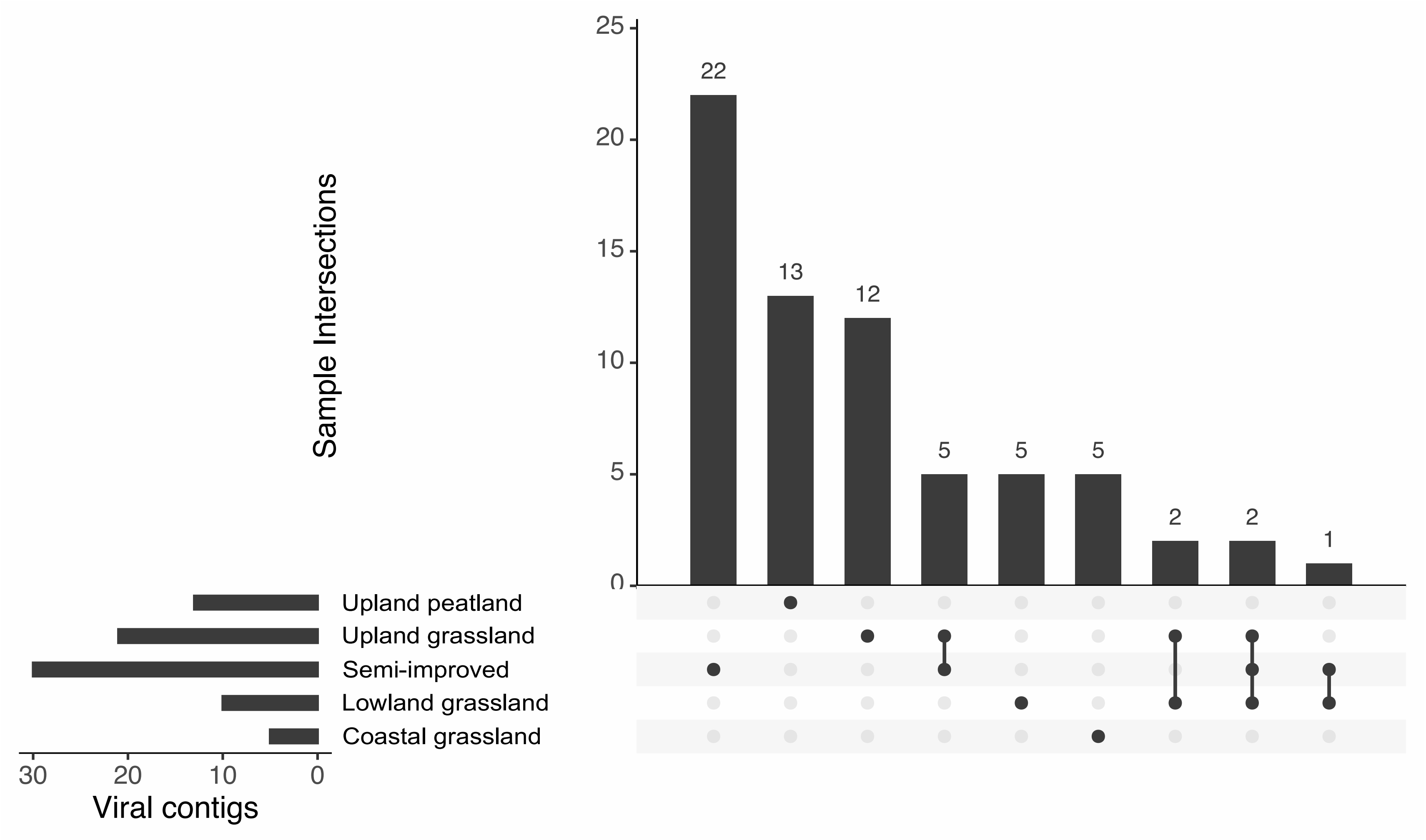


**Supplementary Figure 4** – Pruned phylogenetic tree (a) and site distribution (b) of putative dicistro-like viruses. Viruses were predominantly found within upland/ semi-improved sites.

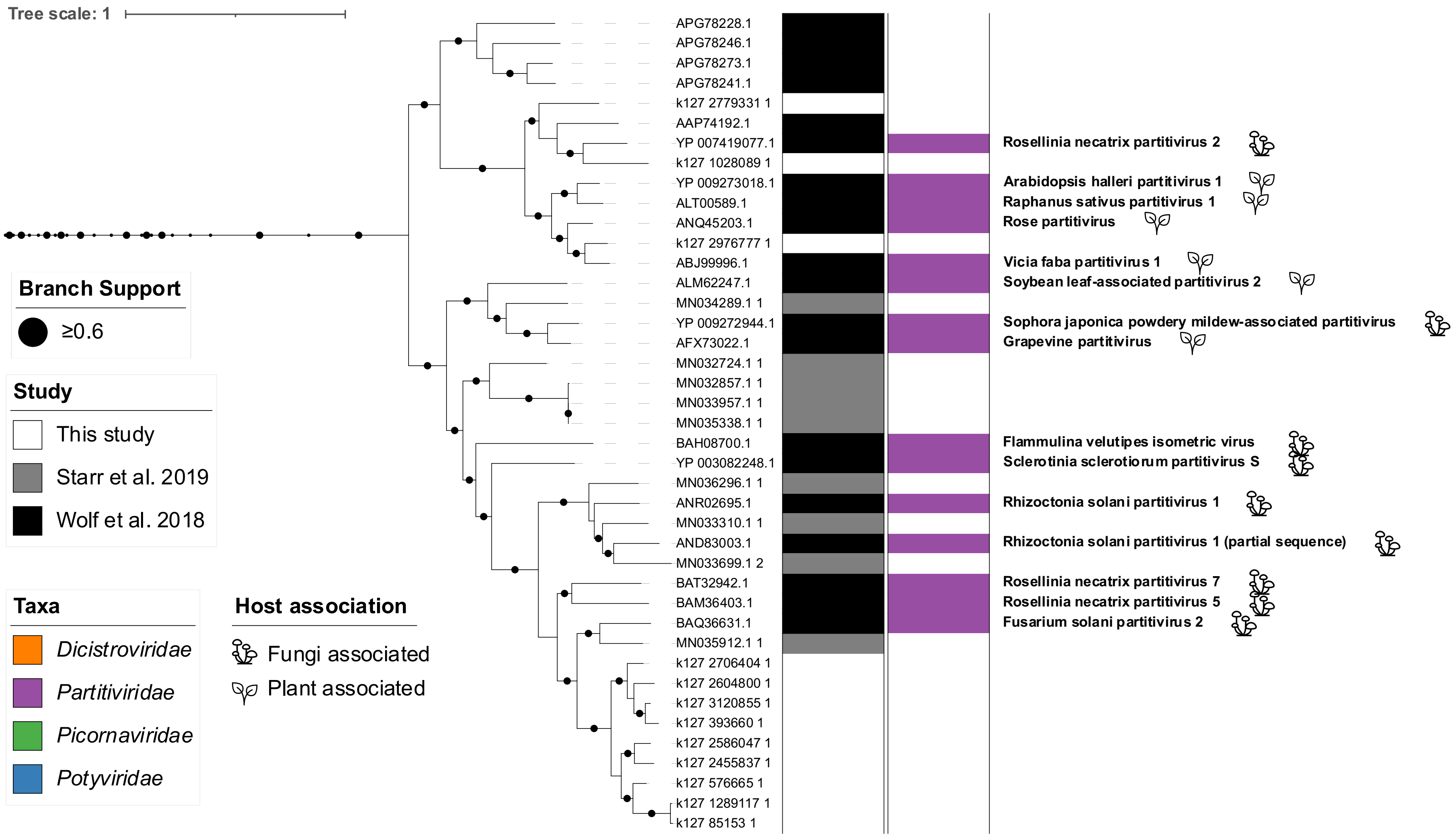


**Supplementary Figure 5** – Pruned phylogenetic tree of putative partiti-like viruses. The majority of partiti-like vOTUs identified in this study are relatively closely related to Fusarium solani partitivirus 2 (indicated by short branch lengths at the bottom of the figure).

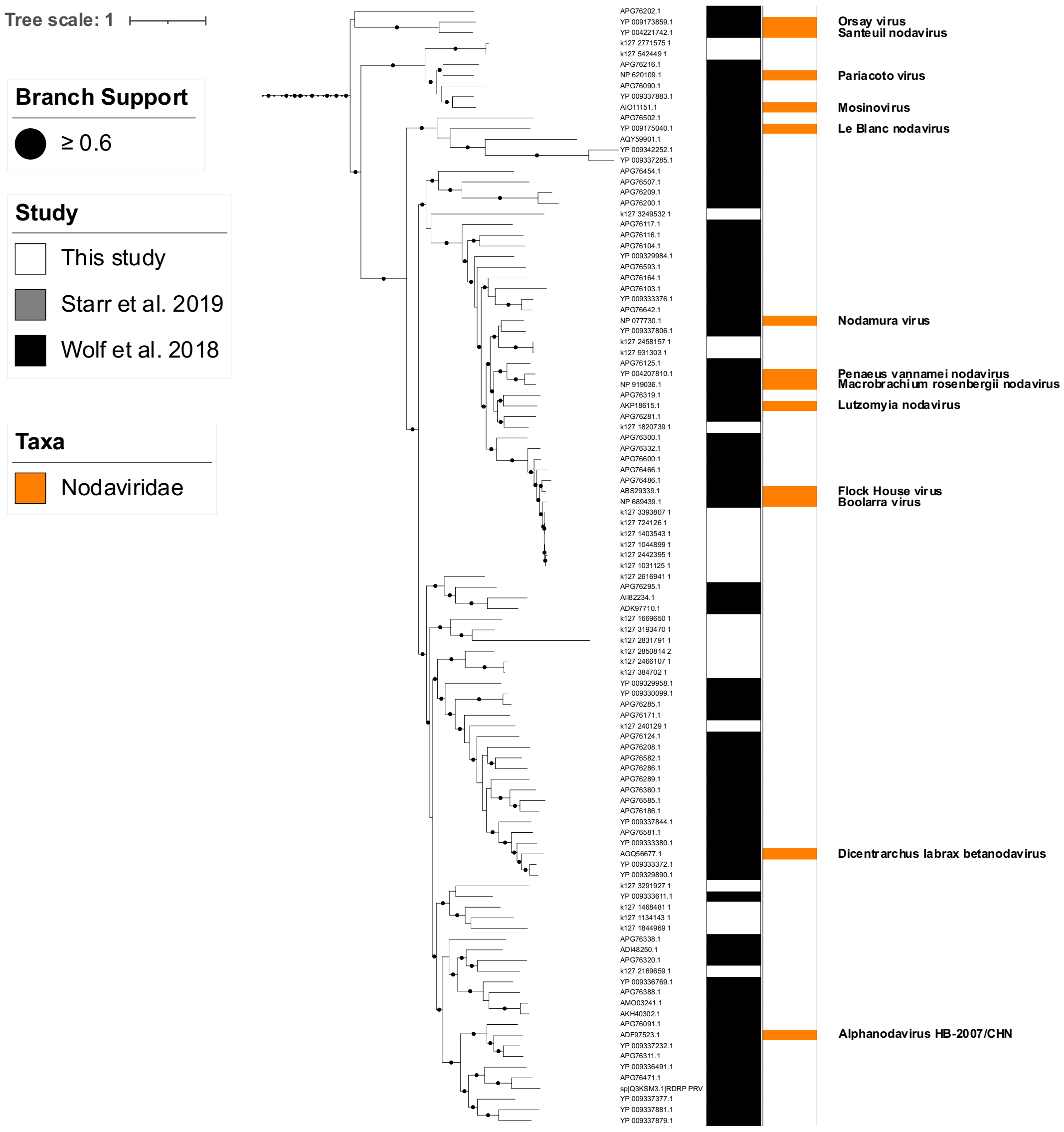
Tree scale: 1
Branch Support
≥ 0.6
Study
This study
Starr et al. 2019
Wolf et al. 2018
Taxa
Nodaviridae
Orsay virus
Santeuil nodavirus
Pariacoto virus
Mosinovirus
Le Blanc nodavirus
Nodamura virus
Penaeus vannamei nodavirus
Macrobrachium rosenbergii nodavirus
Lutzomyia nodavirus
Flock House virus
Boolarra virus
Dicentrarchus labrax betanodavirus
Alphanodavirus HB-2007/CHN

(a)

(b)

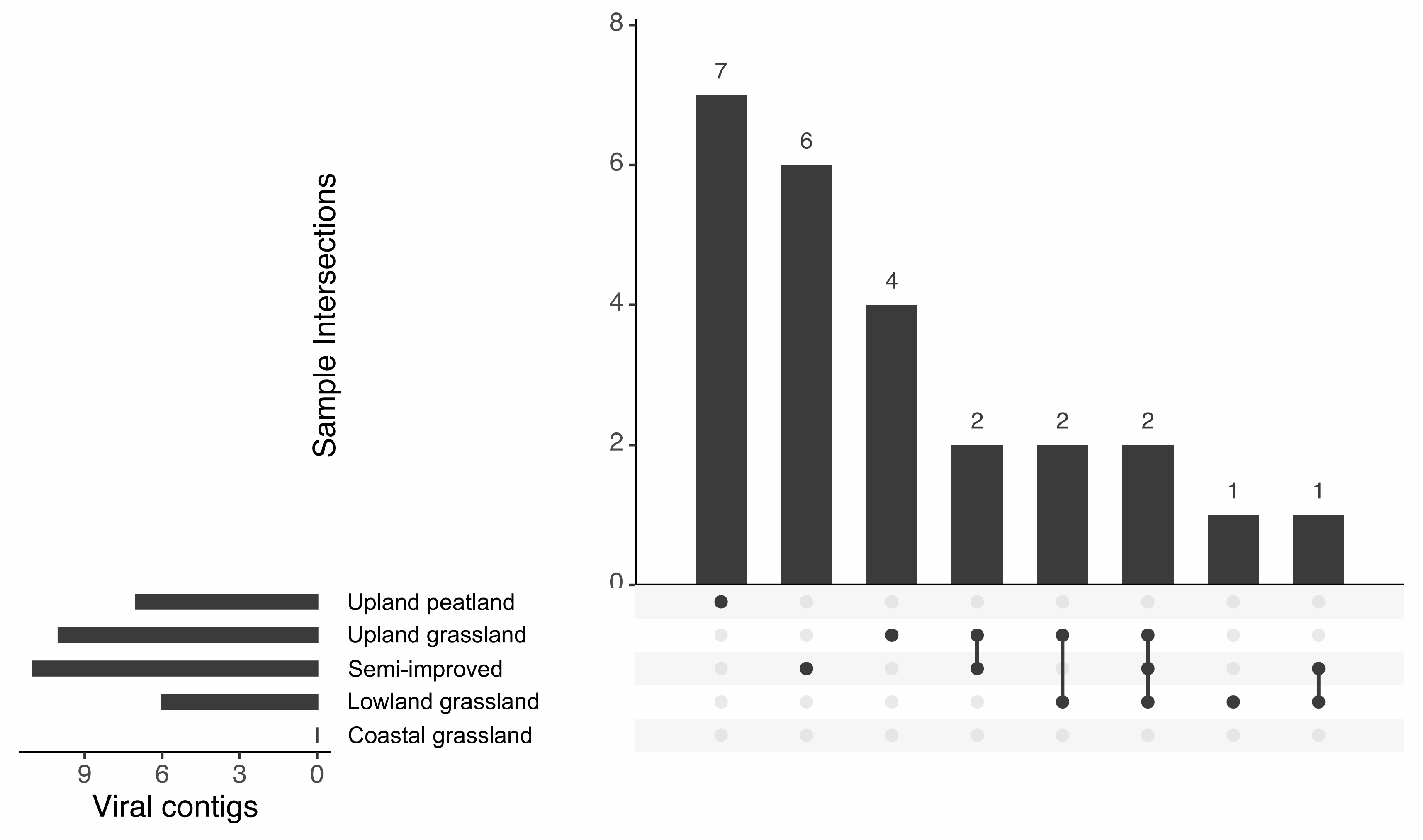


**Supplementary Figure 6** – Enlarged phylogenetic tree (a) and site distribution (b) of noda-like viruses found in this study clustering near reference *Nodaviridae* RdRP sequences (a). Viral sequences were predominantly found in upland areas, with no noda-like viruses found in the coastal grassland site.
